# Fragment-based *ab initio* phasing of peptidic nanocrystals by MicroED

**DOI:** 10.1101/2021.09.13.459692

**Authors:** Logan S. Richards, Maria D. Flores, Claudia Millán, Calina Glynn, Chih-Te Zee, Michael R. Sawaya, Marcus Gallagher-Jones, Rafael J. Borges, Isabel Usón, Jose A. Rodriguez

**Affiliations:** Department of Chemistry and Biochemistry; UCLA-DOE Institute for Genomics and Proteomics; STROBE, NSF Science and Technology Center; University of California, Los Angeles (UCLA); Los Angeles, CA 90095, USA; Crystallographic Methods, Institute of Molecular Biology of Barcelona (IBMB–CSIC), Barcelona Science Park, Helix Building, Baldiri Reixach 15, 08028 Barcelona, Spain; Department of Biological Chemistry and Department of Chemistry and Biochemistry, University of California Los Angeles (UCLA), Howard Hughes Medical Institute (HHMI), UCLA-DOE Institute for Genomics and Proteomics, Los Angeles, CA 90095, USA; ICREA, Institució Catalana de Recerca i Estudis Avançats, Passeig Lluís Companys 23, 08003 Barcelona, Spain

**Keywords:** Phasing, *Ab initio*, Fragment-based phasing, MicroED, Nanocrystal, Peptide, Cryo-EM, ARCIMBOLDO

## Abstract

Microcrystal electron diffraction (MicroED) is transforming the visualization of molecules from nanocrystals, rendering their three-dimensional atomic structures from previously unamenable samples. Peptidic structures determined by MicroED include naturally occurring peptides, synthetic protein fragments and peptide-based natural products. However, as a diffraction method, MicroED is beholden to the phase problem, and its *de novo* determination of structures remains a challenge. ARCIMBOLDO, an automated, fragment-based approach to structure determination. It eliminates the need for atomic resolution, instead enforcing stereochemical constraints through libraries of small model fragments, and discerning congruent motifs in solution space to ensure validation. This approach expands the reach of MicroED to presently inaccessible peptidic structures including segments of human amyloids, and yeast and mammalian prions, and portends a more general phasing solution while limiting model bias for a wider set of chemical structures.

## Introduction

Crystallography has played a momentous role in our understanding of peptidic structures^1^. Micro electron diffraction (MicroED) is expanding its scope by delivering atomic structures from peptide crystals less than a micron in thickness^2–4^. Electron diffraction leverages the strong interaction of electrons with matter, capturing diffraction signal that would be missed by conventional X-ray crystallography^5^. Some molecules of high biological or chemical importance are only known to grow nanocrystals, demanding structural methods of extreme sensitivity, as seen in the amyloid peptide structures of the toxic core of the Parkinson’s-associated protein α-synuclein^3^ or the ultra-high-resolution structure of a prion protofibril^4^. Likewise, the technique has determined structures of complex bio-derived or post-translationally modified peptides such as an amyloid-β core with a racemized residue^6^, the cyclic peptide antibiotic thiostrepton^7^ and a synthetic tetrapeptide natural product analogue^8^.

Current determination of MicroED structures follows one of two routes: *ab initio* phasing through direct methods^9^ if data resolution is atomic, or molecular replacement (MR)^10^ when a highly similar structure is known^3,11–13^. MR is challenged by unknown peptide structures that contain uncharacterized backbone geometries or a substantial fraction of unnatural amino acids. Without atomic resolution data, novel phasing solutions are still needed for MicroED targets of uncertain geometry, identity, or chemical connectivity.

Fragment-based phasing (FBP)^14^ yields accurate solutions relying on the computational search for defined subsets of a target structure to obviate the need for atomic resolution data. Fragments are located by likelihood-based molecular replacement^15^ and expanded through density modification and map interpretation^16^. The ARCIMBOLDO programs substitute the atomicity constraint underlying direct methods with stereochemical constraints^17^. For a structure containing defined fragments of constant geometry a single model fragment is appropriate and model alpha-helices have been particularly successful^18^. General cases require one jointly evaluate libraries of fragments, representing variations of a structural hypothesis. Relying on secondary and tertiary structure fragments extracted from the Protein Data Bank (PDB)^19^ or from distant homologs^20^, ARCIMBOLDO_BORGES has been broadly used in phasing protein structures determined by X-ray crystallography. ARCIMBOLDO has also been used on MicroED data, to phase a 1.6Å of Proteinase K from distant homologues^21^. Where fragments are placed accurately, they can yield solutions despite accounting for as little as 4% of the scattering atoms in a structure. Since any experimental or calculated fragment may be used as input, fragment-based phasing could prove powerful for the general determination of peptidic or other chemical structures by electron diffraction.

Here, we showcase fragment-based phasing for *ab initio* structure determination of novel peptide structures from MicroED data in the absence of atomic resolution. Our approach is based on the development of new library methods tailored to sample structural variability, while profiting from the reduced size of active peptide structures to preclude model bias. We validate its success on known and novel structures obtained from nanocrystallites formed by diverse amyloid peptides.

## Results

### High-resolution MicroED data from peptide nanocrystals

The limited crystal size and directional growth exhibited by some peptides of high biological or chemical interest renders electron diffraction a necessary choice for structure determination. However, faced with a growing number of MicroED datasets from peptide crystals, for which direct methods and molecular replacement solutions were unavailable, we set out to develop dedicated fragment phasing approaches for this set of substrates.

Nanocrystals from each of five peptide segments summarized in Table 1 were preserved on grids in a frozen-hydrated state; crystals of each were visually identified and diffracted as previously described^3^. Ideal candidates for MicroED yielded better than 2Å diffraction. Diffraction data from several crystals were merged to improve completeness. Structural determination via direct methods with SHELXD^9^ succeeded for peptides whose crystals diffracted to atomic resolution: a synthetic mammalian prion segment (QYNNENNFV) **[1]** and a sequence variant of a repeat segment of the yeast prion New1p (NYNNYQ) **[2]** as well as a plate polymorph of the functional mammalian prion, CPEB3 (QIGLAQTQ) **[3]**. Direct methods solutions were unattainable for the needle polymorph of **[3]**, a segment of the human amyloid protein LECT2 (GSTVYAPFT) **[4]** and a segment from the human zinc finger protein (ZFP) 292 (FRNWQAYMQ) **[5]**.

**Table 1:**
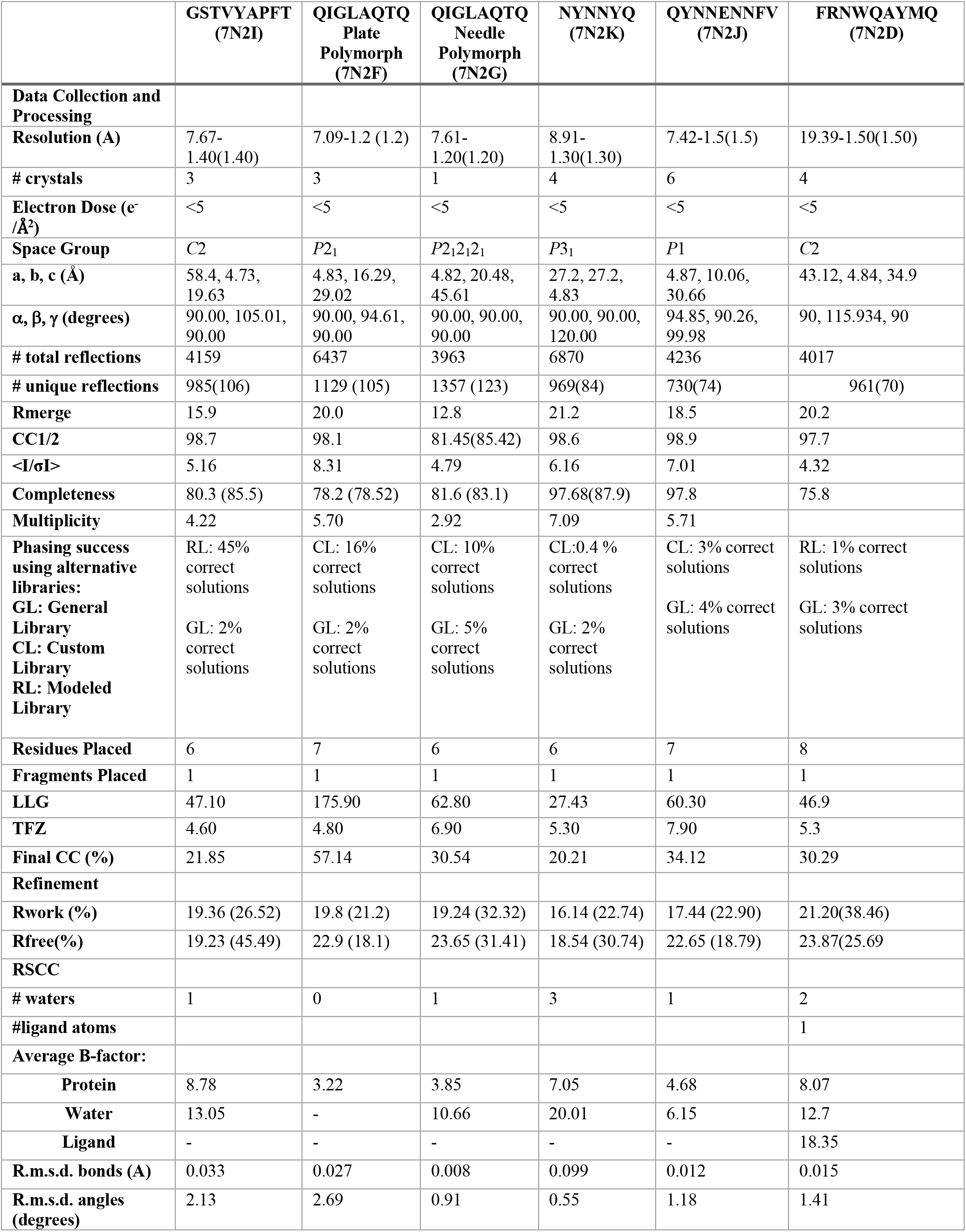
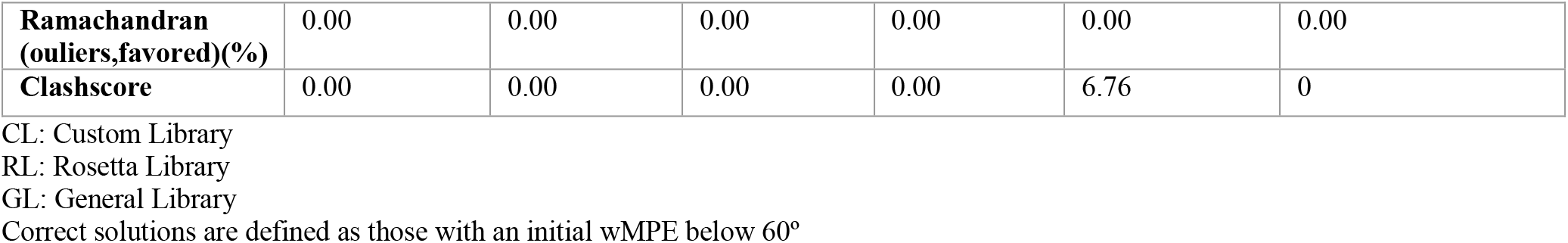

### Phasing with ARCIMBOLDO using general local fold libraries

For *ab initio* macromolecular phasing, ARCIMBOLDO_BORGES exploits fragment libraries representing a common local fold as found in a vast number of PDB structures. Such libraries can be derived from millions of fragments clustered to describe the geometrical variation within the radius of convergence of the method. Fragment superposition allows joint statistical analysis of all phasing attempts as a single experiment. Since typical fragments in macromolecular libraries contain more residues than our peptide structures^22^, we devised dedicated libraries that handled the extreme abundance of motifs exhibited by short peptides; for example, two short antiparallel beta strands. Weighting overall and local properties when superposing such small models also presented a challenge that was solved experimentally, simulating data from a template and refining the location of all library models against the calculated data.

Single-strand libraries yielded partial solutions for all peptides in Table 1. However, the best solutions identified by this approach did not benefit from the statistical or phase combination algorithms enabled by prior superposition^23^. Instead, optimal solutions were obtained by correct placement of the single most accurate fragments. As peptidic MicroED datasets tend to be small and thus amenable to a large number of calculations from different starting fragments, we decided to exploit competing hypotheses to address the more pressing concern of model bias. With that in mind, we devised libraries encompassing dissimilar, non-superimposable models.

### Broad, knowledge-based libraries to phase nanocrystals

To develop diverse, knowledge-based libraries, we began with ideal cases: atomic resolution data from crystals of two peptide segments that yielded structures by direct methods. The first dataset combined data from five crystals of a mammalian prion segment **[1]**. The full 0.9Å data were included in our set as a gold standard (Figure 1A). When intentionally truncated to a resolution of 1.5 Å, both direct methods and molecular replacement using models of closely related prion sequences^4^ were unsuccessful. A library of 249 poly-glycine hexapeptides derived from previously determined amyloid peptide structures was used within ARCIMBOLDO_BORGES (Figure 1B). This library produced clearly discriminated potential solutions scoring above 25% in their initial correlation coefficient^24^ (Figure 1C). These same solutions would later be found to exhibit low errors relative to the phases calculated from a final structure (Table S2). The best solution placed six alanine residues and was sufficient to build the remainder of the peptide based on difference density (Table 1, Figure 1D/E).

**Figure 1:**
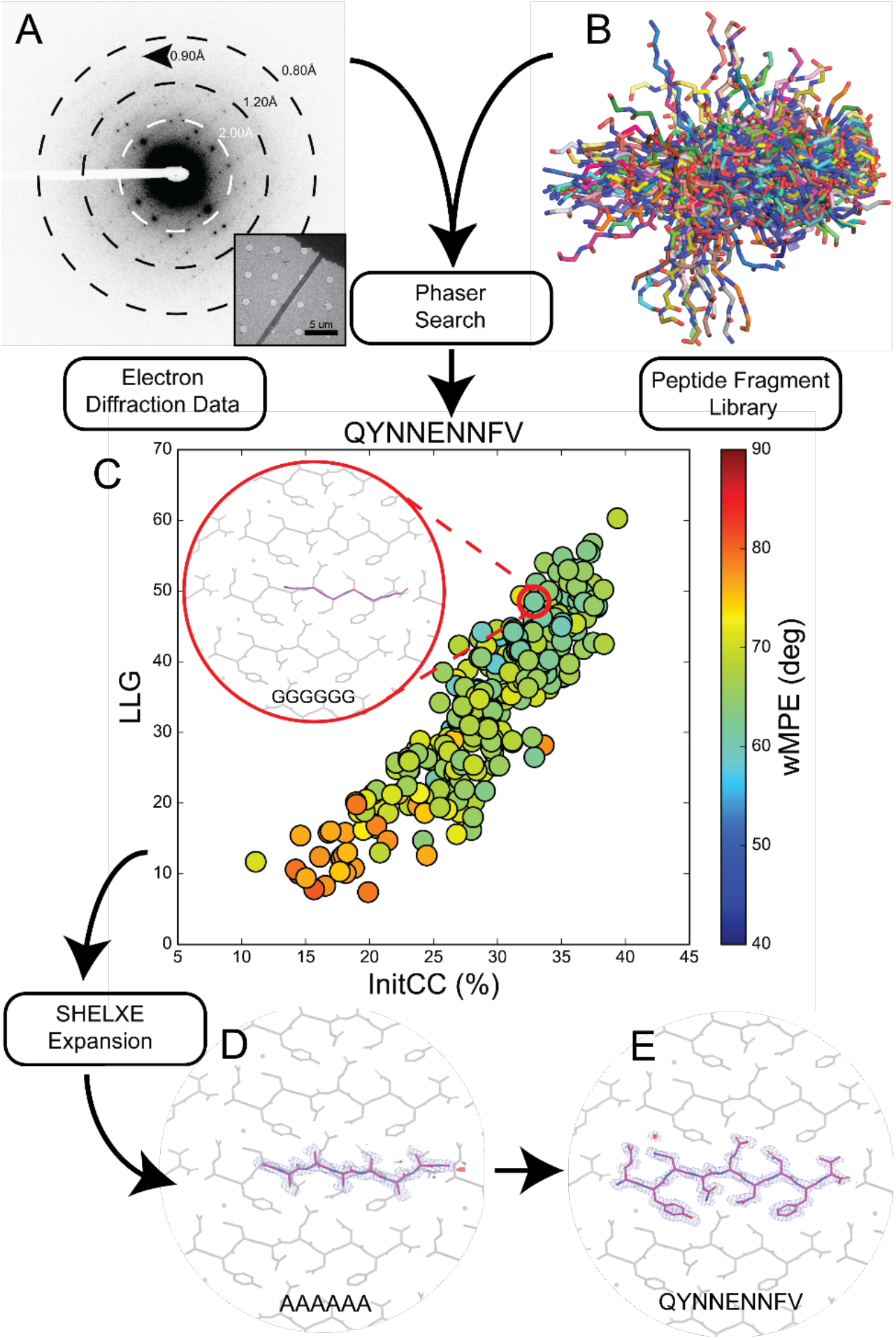
Workflow for using peptide fragments for phasing MicroED data. A) Electron diffraction pattern reaching atomic resolution for QYNNENNFV. Rings designate resolution ranges while arrowhead designates highest resolution spot. B) All fragments comprising the poly-glycine hexapeptide library aligned in pymol. C) LLG vs. InitCC plot for the fragments screened in ARCIMBOLDO-BORGES. Color indicates the wMPE of the fragment relative to the phases calculated from the final structure. Inset shows the fragment chosen for SHELXE expansion overlaid on the final structure. D) The output solution from ARCIMBOLDO-BORGES following SHELXE expansion is shown overlaid on the final structure. Maps are shown after one round of refinement in Phenix. E) Final structure and potential map for the QYNNENNFV peptide with symmetry related chains shown in grey

Data collected from crystals of the New1p segment **[2]** presented an increased challenge for FBP. Although microcrystals of **[2]** yielded an X-ray crystallographic structure (Figure S2, S3B), the same condition also produced nanocrystals requiring MicroED. The 1.1Å dataset obtained combining 4 crystals of this polymorph rendered by direct methods a different structure from that originally determined by X-ray diffraction (Figure S3A/B). The dataset truncated to 1.3Å served as a second test for ARCIMBOLDO_BORGES. A library holding poly-glycine pentapeptides yielded a single promising solution that could be fully extended and matched the direct methods solution (Table 1, Figure S4).

### Structure from a segment of the prion domain of CPEB3 [3]

We next sought to determine novel peptide structures from data outside the reach of direct methods. Crystallization of a segment from the prion domain of CPEB3 **[3]** produced crystal slurries suitable for MicroED (Figure 2). Screening them in overfocused diffraction mode revealed two distinct morphologies, suggesting the presence of multiple structures (Figure 2A/E). Crystals of plate morphology belonged to *P*2_1_, while those of the relatively rare needle morphology presented space group *P*2_1_2_1_2_1_ (Table 1, Figure 2E).

**Figure 2:**
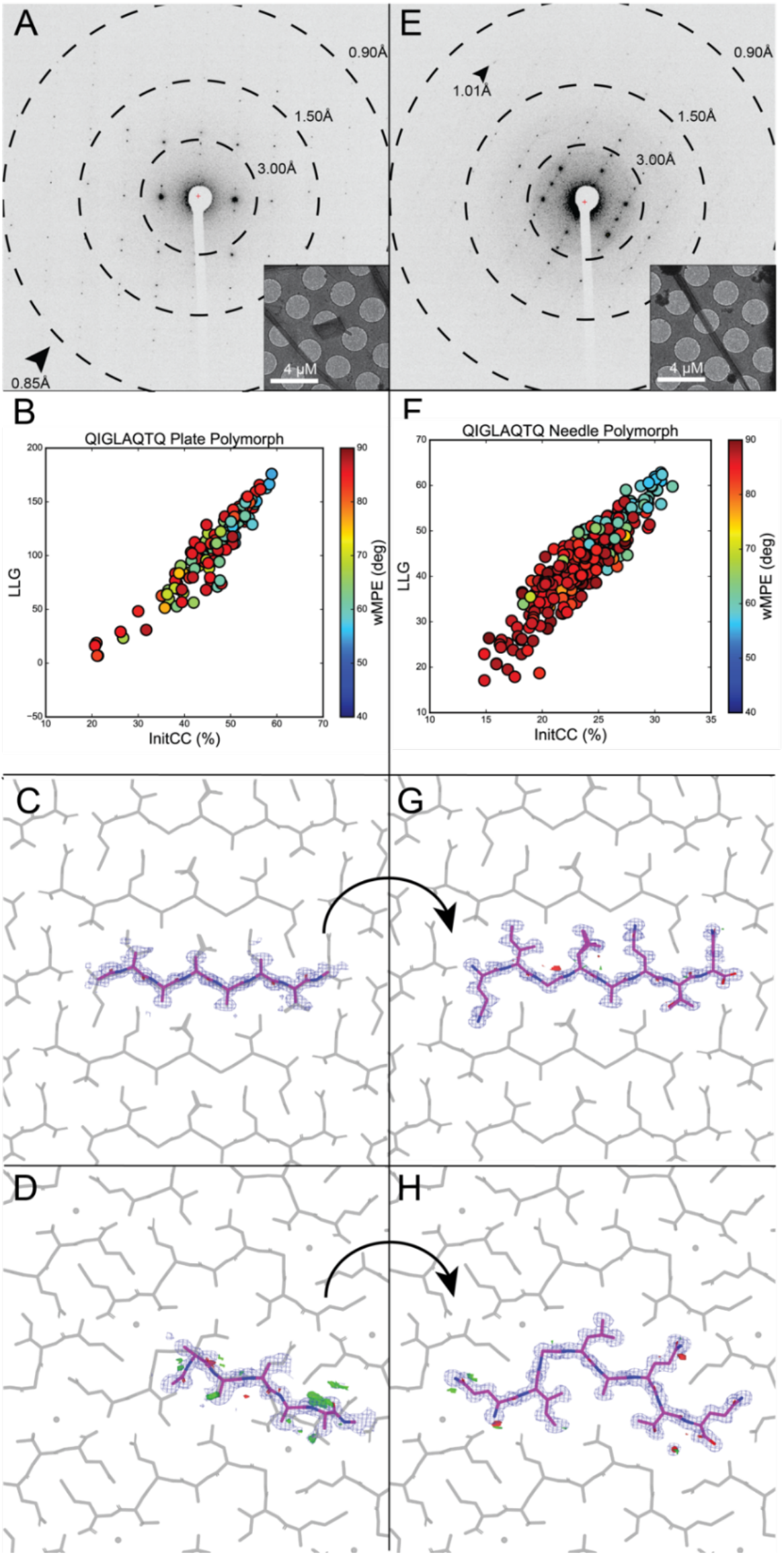
FBP of MicroED data from CPEB3 peptide QIGLAQTQ reveals two polymorphs. A) Electron diffraction pattern reaching atomic resolution for QIGLAQTQ plate morphology. Rings designate resolution ranges while arrowheads designate the highest resolution spot. Overfocused diffraction image of plate morphology crystal. B) Post-mortem analysis from ARCIMBOLDO-BORGES plotting LLG vs. InitCC (*P*2_1_). C) Initial output potential maps following SHELXE expansion by ARCIMBOLDO-BORGES for plate polymorph overlaid on final solution (grey). Buildable density visible on several residues. D) Final density maps of QIGLAQTQ plate polymorph asymmetric unit with symmetry mates shown in grey. E) Diffraction pattern for QIGLAQTQ needle morphology. Rings designate resolution ranges while arrowhead designates highest resolution spot. F) Post-mortem analysis from ARCIMBOLDO-BORGES plotting LLG vs. IniCC (P 2_1_ 2_1_ 2_1_). G) Initial output potential maps following SHELXE expansion by ARCIMBOLDO-BORGES for needle polymorph overlaid on final solution (grey). H) Final density maps of QIGLAQTQ needle polymorph asymmetric unit with symmetry mates shown in grey and water molecule displayed as red sphere.

Merged data from 3 crystals of the plate polymorph were phased by direct methods. Its structure, an unkinked beta strand, was refined at 1.0Å (Figure S3C). In contrast, a single crystal of the rare needle polymorph diffracted to ∼1.1Å resolution, generated a dataset that was 81.5% complete at 1.2Å (Figure 2). Despite its high resolution, neither SHELXD nor molecular replacement with the structure derived from the plate polymorph truncated to poly-alanine yielded a solution. Instead, the needle polymorph dataset was successfully phased using a 270-fragment poly-glycine library of tetrapeptides in ARCIMBOLDO-BORGES (Table 1, Figure S1A/B). An initial solution containing four alanine residues led to a fully refined model (Figure 2G/H). Alternatively, applying the same procedure with an 89-fragment poly-glycine library of pentapeptides (Figure S1C/D) to data from the plate polymorph also resulted in a number of possible solutions. In both cases, non-random solutions with the highest LLG and Initial CC scores were identified by ARCIMBOLDO_BORGES (Table S2, Figure 2B/F).

### Structures determined by modelled fragment libraries

To overcome the limitation of requiring prior structural knowledge for fragment libraries, we generated atomic models computationally. Such libraries would be ideally suited for determining structures with unanticipated local geometries and could be broadly applicable to small molecules. We computed fragments using PyRosetta^25,26^ starting from a known peptide backbone as a template onto which sequences of interest were threaded and modelled^27^ (Figure 3). This scheme was parallelized to generate libraries containing hundreds of fragments, which successfully facilitated FBP of several unknown structures:

**Figure 3:**
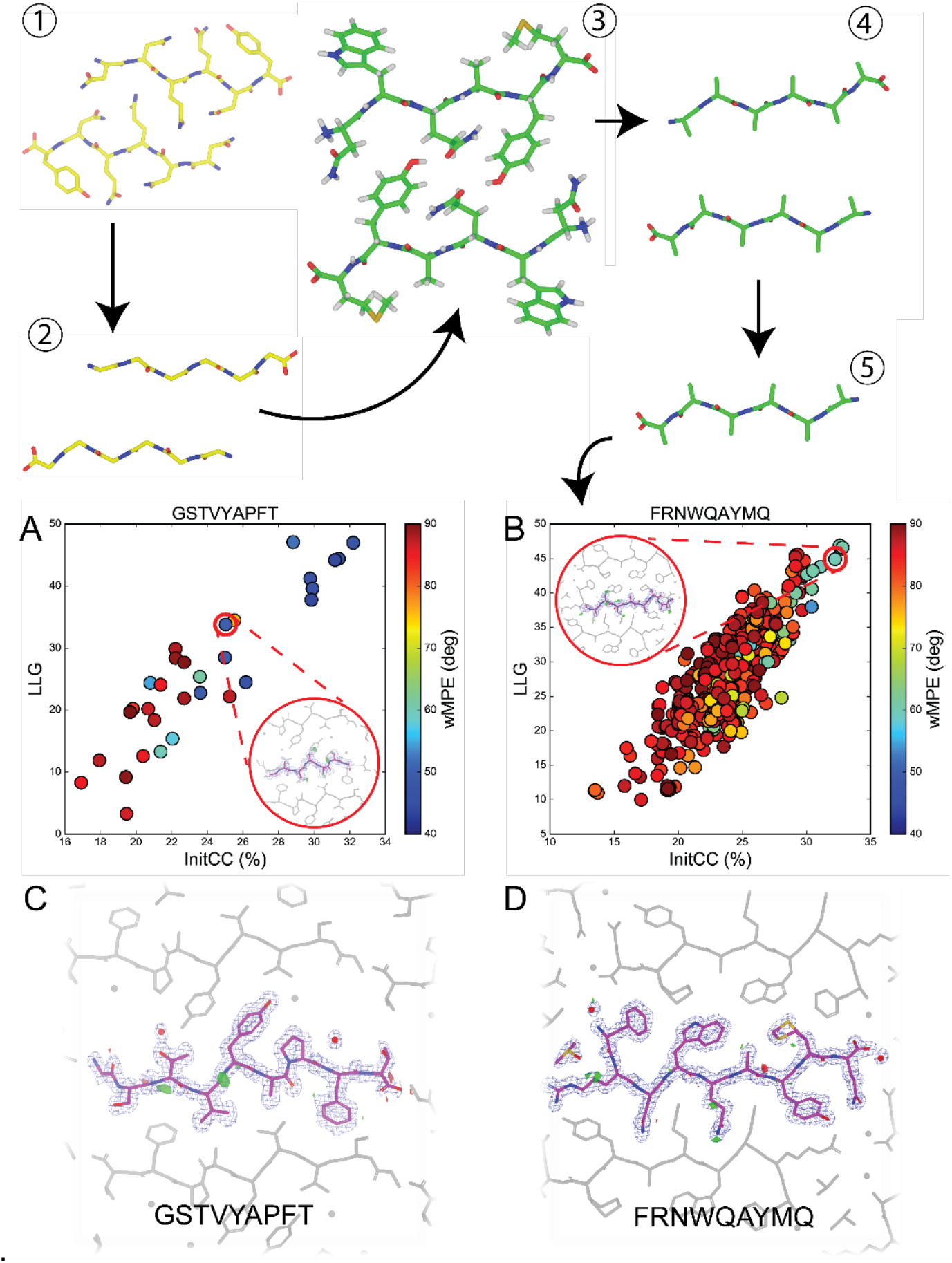
Rosetta library generation and wMPE analysis for GSTVYAPFT and FRNWQAYMQ peptide structures. 1) The NNQQNY peptide structure used as a template steric zipper for Rosetta modelling. 2) The NNQQNY peptide stripped to glycine residues in preparation for threading. 3) One example of a Rosetta generated steric zipper structure after threading, repacking, and relaxing in iterative cycles to reach a calculated energy minimum. 4) The Rosetta generated structures are stripped to alanine residues and have hydrogen atoms removed. 5) Individual chains are isolated and are used as the fragment library for phasing with ARCIMBOLDO-BORGES. The LLG vs. InitCC plots for (A) GSTVYAPFT and (B) FRNWQAYMQ are shown below. Inset shows the fragment that was chosen for SHELXE expansion, leading to the correct solution, overlaid with the final structures. C) Final potential maps of GSTVYAPFT asymmetric unit with symmetry mates shown in grey. D) Final potential maps of FRNWQAYMQ asymmetric unit with symmetry mates shown in grey.

Crystals grown from a segment of the LECT2 protein **[4]** diffracted to only 1.4Å by MicroED, and the dataset combined from 3 such crystals failed to yield solutions from direct methods or molecular replacement with full peptide structures (Table 1). We generated a new hexapeptide poly-alanine library using Rosetta in an attempt to approximate the structure of **[4]** in close packing (Figure S1G/H). The full sequence of the LECT2 protein was used to generate hexapeptide models, that were subsequently threaded pairwise onto the two backbones of the known structure of the peptide NNQQNY^27^. Models were allowed to repack and relax to energy minima in Rosetta. Then, 111 models containing the GSTVYAPFT sequence were truncated to alanine and used in ARCIMBOLDO_BORGES (Figure 3), yielding a discriminated solution (Table S2).

A segment from the human protein ZFP-292 **[5]** presented the most severe challenge to conventional phasing. Due to a high degree of orientation bias of crystals on EM grids, merging data from 4 crystals only achieved an overall completeness of 75.8% at 1.5Å resolution (Table 1). As in the case of **[4]**, this segment could not be phased by direct methods or standard molecular replacement, and no solutions were found when attempting FBP using our fragment libraries from known structures. We again turned to Rosetta in this case to populate a library of fragments that approximated the structure of **[5]**, relying only on the 9-residue sequence of the peptide to generate paired hexapeptide sequences for threading. These threaded segments were then evaluated in Rosetta to generate 20 models per pair (Figure 3). This library of 640 models produced in ARCIMBOLDO_BORGES (Figure S1) yielded low wMPE solutions, one of which was refined to completion (Figure 3B/D, Table S2).

## Discussion

To satisfy the need for new *ab initio* phasing solutions for MicroED, we develop and deploy new fragment-based phasing strategies using ARCIMBOLDO_BORGES and determine five novel structures. These include structures that could not be determined by conventional phasing methods. The variation in diffraction quality obtained from these peptides is representative of the spectrum typically encountered in chemical structure analysis, including examples of relatively low completeness, crystals with low solvent content, and lack of atomic resolution. While the six structures determined here represent a small sampling of the greater universe of peptidic molecules, each of these structures revealed challenges that could be generalized. All analyzed peptides had a high aggregation propensity, contained little to no disordered solvent and naturally produced nanocrystallites instead of larger crystals. Polymorphism was encountered and in one case revealed differences in atomic structure.

These issues compromise the unambiguous identification of correct structure solutions, particularly where atomic resolution is not available and information from pre-existing solutions must be used in the phasing process. Precluding errors derived from model bias becomes even more pressing in such cases. Hence, ARCIMBOLDO jointly evaluates large libraries, where competing hypotheses are compared to provide a safeguard against erroneous solutions. The fragments used in our approach were successfully selected by Phaser, based on their LLG, identified with SHELXE CC scores and subsequently expanded by SHELXE into accurate initial solutions. While we observe examples of high phase error models scoring well in preliminary steps, the discrimination of competing potential solutions revealed an unambiguous solution in all cases.

Verification through competition^28^ is particularly promising in chemical crystallography given that exhaustive searches of solution space are manageable. The libraries exploited for macromolecular phasing hold variations around a common local fold and yielded correct solutions in all instances. In all cases, single fragment solutions were found to outperform combined solutions. Broadening the base of hypotheses through the use of heterogeneous libraries was introduced to address model bias. All solutions were verified through clear discrimination between conflicting models and similarly high scores for structurally compatible hypotheses. Knowledge-based and modeled libraries rendered higher Z-scores than general libraries, discriminating as illustrated in Fig2B, F. This makes the present strategy amenable to exploring distinct secondary structure motifs, including primarily ɑ-helical peptides or structures with more than one type of secondary structure^29,30^.

Extending fragment-based phasing to make use of computed fragments broadens the scope of targets for MicroED to unprecedented structures beyond peptidic compounds. Computed libraries succeeded when applied to our most challenging cases [4-5] and could further benefit from new advancements in machine learning. Recently, AlphaFold harnessed the vast structural diversity available in the PDB using deep neural networks to achieve correct prediction of protein folds with unexpectedly high accuracy^31^. However, small, chemically and geometrically diverse structures still require dedicated development. Exploration of the rich structural expanse of chemical space will require methods that accurately select structural fragments while excluding bias artifacts to achieve structural solutions by MicroED.

## Summary

We expand the *ab initio* phasing toolkit for MicroED overcoming the need of atomic resolution diffraction to produce *de novo* solutions. Using ARCIMBOLDO-BORGES and libraries of both known and computed structures, we determine six novel atomic structures of peptide segments. The structures determined using this method are accurate and represent varied geometries and sequences. Model bias is precluded by parallel assessment of a large collection of structural hypotheses providing a baseline. These methods successfully establish a three-dimensional structure from samples that were previously intractable and open a road to structural solutions for small molecules from near atomic-resolution MicroED data.

## Methods

### Collection of microfocus X-ray data and structure determination

Crystal clusters of NYNNYQ were grown at room temperature in a 96-well Wizard screen, using a nominally 24.5 mM aqueous solution of the peptide. The crystallization condition chosen for further optimization in 24-well, hanging drop trays consists of 20% 2-methyl-2,4-pentanediol (MPD), buffered by 0.1M sodium acetate pH 4.5). The peptide crystals were harvested from hanging drops using CryoLoops™ from Hampton Research with no additional cryoprotectant other than the MPD already present and flash-frozen in liquid nitrogen. Diffraction datasets were collected under cryogenic conditions (100 K) on beamline 24-ID-E at the Advanced Photon Source (APS) equipped with an ADSC Q315 CCD detector, using a 5 µm beam with a wavelength of 0.979 Å. The data were collected via manual vector scanning. 56 diffraction images were collected over three scans from one crystal and one scan from a different crystal. All images have an oscillation range of 5° and were indexed and integrated by XDS^32^. The reflection list outputted by XDS was sorted and merged in XSCALE. SHELXD^33^ was able to reach an *ab initio* solution. The atomic coordinates from SHELXD were used to generate a F_calc_ map with SHELXL^34^. An atomic model commensurate with the generated electron density was built in Coot and refined in PHENIX against measured data. The refinement statistics of the final structure are listed in Table S1.

### Preparation of peptide nanocrystals

Lyophilized, synthetic peptides were purchased from Genscript. Crystals of each peptide were grown as follows: the QYNNENNFV peptide was dissolved in water at 0.88 mM. Crystals were grown using the hanging drop method where 1.5 µL of peptide was added to 1.5 µL of well solution - 0.1M Li_2_SO_4_, 2.5M NaCl, 0.1M NaOAc adjusted to pH 4.5 with acetic acid - in 24 well trays over 500 µL of well solution. QIGLAQTQ was prepared at 64.5 mM in water. Crystals were grown via 24-well hanging-drop vapor diffusion at 27 mM in 14% polyethylene glycol (PEG) 20,000, 100mM MES and 3% DMSO. The crystal slurry was briefly sonicated and 0.5 µL was used to seed a new 3 µL drop and repeated three times. FRNWQAYMQ peptide from ZFP-292 was prepared by dissolving at 1.61 mM in water and 3% DMSO. Crystals were grown in batch containing 35% MPD, 100mM MES and 200mM Li_2_SO_4_, in 1:1 ratio of peptide to buffer. GSTVYAPFT peptide was prepared at 21.3 mM concentration dissolved in water. Crystals were grown in a 10 uL batch in 0.7M sodium formate with 100mM sodium acetate pH 4.6. All crystals appeared within 24 hours and were identified by light microscopy, and subsequently, EM. Batch crystals of NYNNYQ were prepared by dissolving the peptide in water, resulting in a 24.5 mM solution. An equal volume of the crystallization reagent (20% MPD, buffered by 0.1M of sodium acetate to pH 4.5) was added to the peptide solution. The solution was then seeded with crushed crystals grown in hanging drop experiments described in the microfocus X-ray data collection section above.

### Collection of MicroED data

For GSTVYAPFT and QYNNENNFV, 2 uL of crystal slurry was applied to each side of a glow-discharged holey carbon grid (Quantifoil, R 1/4 300 mesh Cu, Electron Microscopy Sciences) followed by plunging into liquid ethane using an FEI Vitrobot Mark IV set to a blot time of 22 and a blot force of 22 for QYNNENNFV and 24 for GSTVYAPFT. For QIGLAQTQ, 1.8 uL of crystal slurry was applied to each side of a glow-discharged holey carbon grid (Quantifoil, R 2/1 200 mesh Cu, Electron Microscopy Sciences) and plunge frozen into liquid ethane using a FEI Vitrobot Mark IV using a blot time of 25-30 seconds and a blot force of 22. For NYNNYQ, 2 uL of crystal slurry was applied to each side of a glow-discharged holey carbon grid (Quantifoil, R 2/1 200 mesh Cu, Electron Microscopy Sciences) and plunge frozen into liquid ethane using a FEI Vitrobot Mark IV, blotting with the force set at 22 for 20-30 seconds.

For GSTVYAPFT, QYNNENNFV, and FRNWQAYMQ diffraction patterns and crystal images were collected under cryogenic conditions using a FEI Tecnai F20 operated at 200keV in diffraction mode. Diffraction patterns were recorded while continuously rotating at 0.3 degrees per second (GSTVYAPFT) or 0.25 degrees per second (QYNNENNFV, FRNWQAYMQ) using a bottom mount TemCam-F416 CMOS camera (TVIPS). Individual image frames were acquired with 2 second exposures per image for all peptides and 5s exposures for some FRNWQAYMQ datasets to increase signal. A selected area aperture corresponding to approximately 4 or 6 μm at the sample plane was selected depending on the crystal. For QIGLAQTQ, diffraction patterns were collected under cryogenic conditions using a Thermo-Fisher Talos Arctica electron microscope operating at 200keV and a Thermo-Fisher CetaD CMOS detector in rolling shutter mode. Individual frames were acquired with 3 second exposures rotating at 0.3 degrees per second using selected area apertures of 100, 150, or 200 microns, as needed to match the size of the crystal. A total of 28 movies were collected from QIGLAQTQ crystals of two distinct morphologies, 15 from crystals of needle morphology and 13 from crystals of plate morphology.

### MicroED data processing

The collected TVIPS movies were converted to individual images in Super Marty View (SMV) format, which are compatible with x-ray data processing software. The diffraction images were indexed and integrated with XDS. The indexing raster size and scan pattern as well as integration in XDS were optimized to minimize contributions by background and intensities from secondary crystal lattices. The reflection outputs from XDS were sorted and merged in XSCALE. For the linear *P*2_1_ QIGLAQTQ structure, 3 partial datasets, containing 252 diffraction images, were merged to produce a final dataset with acceptable completeness (∼80%) up to 1.00 Å, which was truncated to 1.2 Å for phasing and refinement with ARCIMBOLDO. For the kinked *P*2_1_ 2_1_ 2_1_ QIGLAQTQ structure, one dataset, consisting of 82 diffraction images, was sufficient to produce a final merged dataset with high completeness up to 1.2Å for phasing and refinement by ARCIMBOLDO. For the NYNNYQ structure, 4 partial datasets, composed of 297 diffraction images, were merged to produce a final dataset with high completeness up to 1.10Å, which was truncated to 1.3 Å for phasing and refinement with ARCIMBOLDO. For the GSTVYAPFT structure, 3 partial datasets, comprised of 327 diffraction images, were merged to produce a final dataset with high completeness up to 1.3 Å. For the QYNNENNFV structure, 6 partial datasets, containing 931 diffraction images, were merged to produce a final dataset with high completeness up to 0.9 Å, which was truncated to 1.5 Å for phasing and refinement with ARCIMBOLDO. For FRNWQAYMQ structure, 4 partial datasets composed of 224 diffraction images were merged to produce a final dataset with acceptable completeness out to 1.5Å. The statistics for each merge are presented in Table 1.

### Structure determination by direct methods and refinement

Electron diffraction data for NYNNYQ, QIGLAQTQ (plate), and QYNNENNFV were of high enough resolution to yield direct methods solutions. SHELXD was able to reach *ab initio* solutions with all three datasets. The atomic coordinates from SHELXD and corresponding reflection files were inputted into SHELXL^34^ to generate calculated density maps for each solution. Atomic models consistent with the generated density maps were built in Coot and refined in PHENIX against measured data, using electron scattering form factors. The refinement statistics of the final structures are listed in Table S1.

### Generation and use of peptide fragment libraries for phasing MicroED peptide structures using ARCIMBOLDO-BORGES

In ARCIMBOLDO, fragments are identified by likelihood-based molecular replacement^15^ and expanded through density modification and map interpretation^16^. A library of amyloid peptide structures determined by both x-ray and electron diffraction was assembled to provide a diverse collection of backbone conformations with potential in phasing novel structures. To take advantage of these probes, a high-throughput, fragment-based phasing methodology in the form of the ARCIMBOLDO software was used. The ARCIMBOLDO suite of programs uses secondary structure fragments as initial probes for molecular replacement carried out by Phaser^35^. These fragments undergo rotation and translation analysis and are scored based on log-likelihood gain (LLG) and an initial correlation coefficient (InitCC) to identify potentially accurate starting models (Figure 1C). Rather than making an arbitrary choice on how to direct the superposition, determining the best average for the whole model or of a core, or tolerating outlier atoms to be excluded, an empirical answer was drawn simulating data from a template and refining the location of all other models against the calculated data.

Following this, initial maps are calculated and improved by density modification using the sphere-of-influence algorithm in SHELXE^16^. Finally, main chain auto-tracing is performed and solutions are scored by correlation coefficient^36^. Generally, a final CC greater than 25% is indicative of a correct solution. The ability to expand partial solutions in SHELXE permits the use of smaller, potentially more accurate, molecular replacement probes that are identified by Phaser. All runs were carried out using ARCIMBOLDO_BORGES version 2020 Phaser version 2.8.3, and the distributed SHELXE version 2019. Electron scattering factors were used for Phaser analysis and information including molecular weight and predicted solvent content were provided in addition to the reflections data in the form of .mtz and .hkl files.

### Generation of custom and Rosetta fragment libraries

The fragment libraries used as probes were generated in multiple ways. Poly-glycine libraries of varying sizes, four to six residues in length, were generated by extracting fragments from a collection of previously solved amyloid peptide structures (supplementary citations). These fragments were separated into custom libraries only containing fragments of the same length, aligned in PyMOL and used as probes in ARCIMBOLDO_BORGES. The peptides QIGLAQTQ (plate) and NYNNYQ were phased using a poly-glycine library of amino acid pentapeptides containing 89 fragments. The peptide QIGLAQTQ (needle) was phased using a poly-glycine library of amino acid tetrapeptides containing 270 fragments. The peptide QYNNENNFV was phased using a poly-glycine library of amino acid hexapeptides containing 249 fragments.

Additionally, two libraries of poly-alanine fragments were generated using the program Rosetta^37^ by modelling the packing of the peptide sequences threaded over the steric zipper structure of a previously determined amyloid peptide, NNQQNY^27^. After simple threading over the backbone, the side chains were packed and the chains were allowed to relax to a calculated energy minimum over iterative cycles. These models were isolated as individual fragments six amino acids long, stripped to poly-alanine side chains, and aligned in PyMOL to complete the Rosetta libraries. The Rosetta library for GSTVYAPFT was generated by threading all possible six amino acid segments of the LECT2 sequence against each other and then extracting only those which modelled the peptide sequence of interest. This library contained 111 fragments. The Rosetta library for FRNWQAYMQ was generated by threading all six amino acid long permutations of the peptide sequence against each other pairwise over the NNQQNY backbone. The threading, packing, and relaxation steps were done 20 separate times for each pair of sequences and the resulting library contained 640 single chain fragments.

### General strand libraries generated with ALEPH

The general libraries with strands generated by ALEPH for use in ARCIMBOLDO_BORGES and distributed in CCP4 are all larger in size than any of the peptides investigated, as they contained a minimum of three strands forming a sheet in parallel or antiparallel arrangement. To obtain libraries of one or two strands, we started from the distributed three antiparallel strand (udu) library. Using a template of only two strands, we extracted all compatible models (around 24000) using ALEPH. Using the template model, we then generated an artificial dataset in space group P1 to a resolution of 2.0Å. As the models come from a standard library already, they were superposed originally based on their three strands and also clustered geometrically. To achieve that, we performed a Phaser rigid body refinement against the simulated data, using an rmsd error tolerance of 1.0 Å. We then selected the top LLG-scoring fragments for each of the geometric clusters and took them as our representatives for the two-strand library (246 models). Of those, outliers that did not superimpose well were removed. Only 108 models remained – each a pair of antiparallel beta strands. Then we generated two libraries of a single strand by extracting into separate pdbs from the parent library. This procedure yielded two general libraries to use in our experiments.

### Structure refinement

All structures were refined using Phenix version 1.16-3549-000^38,39^ and Coot version 0.8.9^40^. All refinements used standard settings and the built-in electron scattering tables in Phenix. For each peptide, the final model output, called the best.pdb, from the ARCIMBOLDO-BORGES run was used directly as the model for the first round of refinement in Phenix. One exception was the QIGLAQTQ plate polymorph structure. In this case, the model from ARCIMBOLDO-BORGES was run through an additional round of Phaser molecular replacement to correct for what appeared to be a translational shift in the chain. This could be due to the extremely tight packing of the chains and the lack of any solvent, ordered or disordered, making initial chain placement difficult.

## Acknowledgements

We thank Duilio Cascio (UCLA) for discussions and helpful analysis and the David Eisenberg laboratory (UCLA) for generous access to all of their amyloid peptide structures. This work was performed as part of STROBE, an NSF Science and Technology Center through Grant DMR-1548924. This work is also supported by DOE Grant DE-FC02-02ER63421 and NIH-NIGMS Grant R35 GM128867 and P41GM136508. L.S.R. is supported by the USPHS National Research Service Award 5T32GM008496. M.D.F. was funded by Eugene V. Cota-Robles Fellowship and Ruth L. Kirschstein NRSA GM007185 and is currently funded by a Whitcome Pre-Doctoral Fellowship and a National Science Foundation Graduate Research Fellowship. C.G. was funded by Ruth L. Kirschstein NRSA GM007185 and is currently funded by Ruth L. Kirschstein Predoctoral Individual NRSA, 1F31 AI143368. R.J.B. received fellowship from FAPESP (16/24191-8 and 17/13485-3). C.M. is grateful to MICINN for her BES-2015-071397 scholarship associated with the Structural Biology Maria de Maeztu Unit of Excellence. This work was supported by grants PGC2018-101370-B-100, and MDM2014-0435-01 (the Spanish Ministry of Economy and Competitiveness) and Generalitat de Catalunya (2017SGR-1192). J.A.R. is supported as a Pew Scholar, a Beckman Young Investigator and a Packard Fellow.

## Contributions

J.A.R. and I.U. directed the research. L.S.R., M.D.F., and C.M. generated fragment libraries and performed phasing with ARCIMBOLDO. L.S.R, M.D.F, C.G., C.Z. and J.A.R. crystallized peptides and prepared samples. C.G., M.G.J. and J.A.R. collected diffraction data. M.D.F., L.S.R., C.Z., C.G., M.R.S., and M.G.J processed diffraction data. C.M., I.U. and R.J.B. developed and evaluated MicroED-specific features in the ARCIMBOLDO framework. All authors helped write and provided critical feedback on the article.

## Data availability

The deposited ARCIMBOLDO structures of peptides are QYNNENNFV (7N2J), NYNNYQ (7N2K), QIGLAQTQ_plate (7N2F), QIGLAQTQ_needle (7N2G), GSTVYAPFT (7N2I), and FRNWQAYMQ (7N2D). The deposited direct methods structures of peptides are QYNNENNFV (7N2L), NYNNYQ (7N2H), QIGLAQTQ_plate (7N2E).

## Supplementary Data

**Table S1.** Crystallographic data collection and refinement statistics. Values in parentheses are for the highest-resolution shell.

| Direct Methods Structures | NYNNYQ (7N2H) | NYNNYQ | QIGLAQTQ<br>Plate Polymorph<br>(7N2E) | QYNNENNfV (7N2L) |
| --- | --- | --- | --- | --- |
| **Data Collection and Processing** |  |  |  |  |
| Molecular Formula | C<sub>35</sub>H<sub>46</sub>N<sub>10</sub>O<sub>13</sub> | C<sub>35</sub>H<sub>46</sub>N<sub>10</sub>O<sub>13</sub> | C<sub>31</sub>H<sub>55</sub>N<sub>9</sub>O<sub>11</sub> | C<sub>49</sub>H<sub>68</sub>N<sub>14</sub>O<sub>18</sub> |
| No. of crystals merged | 2 (4 Scans) | 4 | 3 | 6 |
| Space Group | P 1 | P 3<sub>1</sub> | P 1 2<sub>1</sub> 1 | P 1 |
| Cell dimensions |  |  |  |  |
| a, b, c (Å) | 4.86, 14.33, 15.75 | 27.20, 27.20, 4.83 | 4.83, 16.29, 29.02 | 4.87, 10.06, 30.66 |
| α, β, γ (°) | 87.369, 86.325, 88.501 | 90, 90, 120 | 90, 94.605, 90 | 94.850, 90.264, 99.988 |
| Resolution Limit (Å) | 1.05 (1.10-1.05) | 1.10 (1.19-1.10) | 1.00 (1.05-1.00) | 0.90 (0.93-0.90) |
| Beam Type | x-ray | electron | electron | electron |
| Wavelength (Å) | 0.979 | 0.0251 | 0.0251 | 0.0251 |
| Electron Dose (e<sup>-</sup>/Å<sup>2</sup>) | N/A | <5 | <5 | <5 |
| Rmerge (%) | 16.1 (35.9) | 21.1 (48.3) | 15.2 (26.2) | 19.7 (47.5) |
| I/σ<sub>1</sub> | 4.13 (1.97) | 4.92 (1.21) | 7.24 (4.03) | 5.21 (2.56) |
| Completeness | 90.2 (61.1) | 83.4 (53.2) | 79.6 (80.8) | 81.0 (82.3) |
| CC<sub>1/2</sub> | 97.4 (89.1) | 98.5 (60.4) | 98.8 (96.8) | 98.8 (78.3) |
| No. total reflections | 4635 (314) | 7893 (431) | 11581 (1430) | 18502 (1603) |
| No. unique reflections | 1793 (154) | 1357 (191) | 1976 (252) | 3437 (330) |
| Multiplicity | 2.6 (2.0) | 5.8 (2.2) | 5.9 (5.7) | 5.4 (4.9) |
| **Refinement** |  |  |  |  |
| Resolution (Å) | 15.70-1.05 (1.10 - 1.05) | 13.60-1.10 (1.19- 1.10) | 7.10-1.00(1.05- 1.00) | 7.98-0.90 (0.93-0.90) |
| No. of Reflections (free) | 1787 (180) | 1356 (136) | 1959 (197) | 3125 (310) |
| Rwork (%) | 14.92 (29.53) | 13.83 (26.25) | 18.60 (22.07) | 20.01 (28.03) |
| Rfree(%) | 16.49 (28.57) | 16.06 (34.89) | 20.92 (25.03) | 20.60 (27.91) |
| CC(work) | 96.9 (0.809) | 98.2 (78.1) | 95.5 (90.3) | 95.8 (81.1) |
| CC(free) | 94.7 (0.679) | 97.2 (52.8) | 92.2 (86.8) | 95.0 (80.5) |
| **Atoms** |  |  |  |  |
| No. peptide atoms | 58 | 58 | 64 | 81 |
| No. waters | 5 | 2 | 0 | 2 |
| No. ligand atoms | 0 | 0 | 0 | 0 |
| Average B-factor: | 5.48 | 7.07 | 3.15 | 4.28 |
| Protein | 4.81 | 6.7 | 3.15 | 4.16 |
| Water | 13.16 | 17.8 | N/A | 8.88 |
| Ligand | N/A | N/A | N/A | N/A |
| R.m.s.d. bonds (Å) | 0.009 | 0.007 | 0.23 | 0.004 |
| R.m.s.d. angles (°) | 0.864 | 0.693 | 1.711 | 0.737 |
| Ramachandran Outliers (%) | 0 | 0 | 0 | 0 |
| Clashscore | 0 | 0 | 0 | 0 |

**Table S2:** Summary of peptide fragment-based phasing

| Peptide | Resolution | # of probes | Best wMPE | Ca RMSD<br>(output to final) | Final R<sub>work</sub>/R<sub>free</sub> |
| --- | --- | --- | --- | --- | --- |
| QYNNENNfV | 1.5Å | 249 | 58.0° | 0.522 Å<sup>2</sup> | 17.44 / 22.65 |
| NYNNYQ | 1.3Å | 89 | 49.5° | 0.561 Å<sup>2</sup> | 16.14 / 18.54 |
| QIGLAQTQ_plate | 1.2Å | 89 | 54.8° | 0.247 Å<sup>2</sup> | 19.8 / 22.9 |
| QIGLAQTQ_needle | 1.2Å | 270 | 55.3° | 2.395 Å<sup>2</sup> | 19.24 / 23.65 |
| GSTVYAPFT | 1.4Å | 111 | 43.9° | 0.509 Å<sup>2</sup> | 19.36 / 19.23 |
| FRNWQAYMQ | 1.5Å | 640 | 53.9° | 1.438 Å<sup>2</sup> | 21.20 / 23.87 |

**Supplementary Figure 1:**
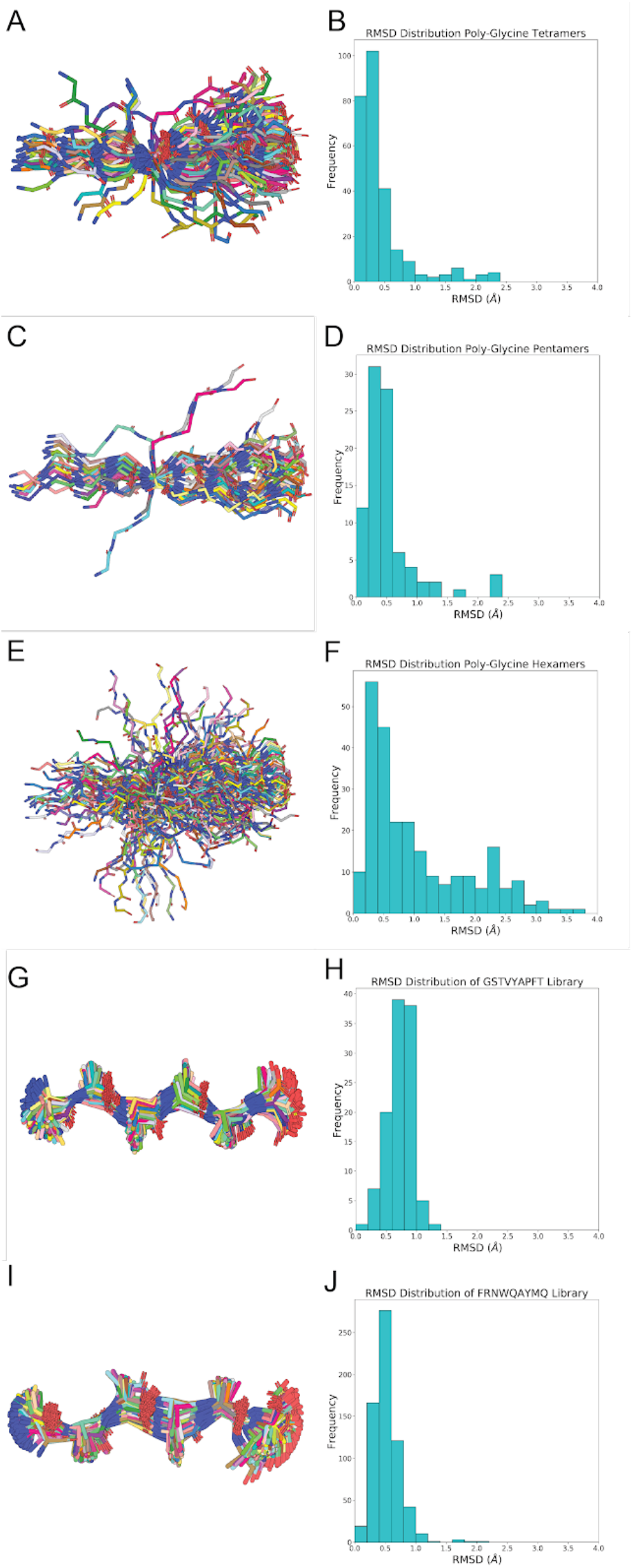
Visualization of fragment libraries and RMSD analysis The poly-glycine tetrapeptide (A), poly-glycine pentapeptide (C), poly-glycine hexapeptide (E), poly-alanine Rosetta GSTVYAPFT (G), and poly-alanine Rosetta FRNWQAYMQ (I) libraries are shown aligned in pymol. B, D, F, H, J) Distribution of RMSD values calculated for each fragment relative to an idealized poly-glycine or poly-alanine strand of equivalent length.

**Supplementary Figure 2:**
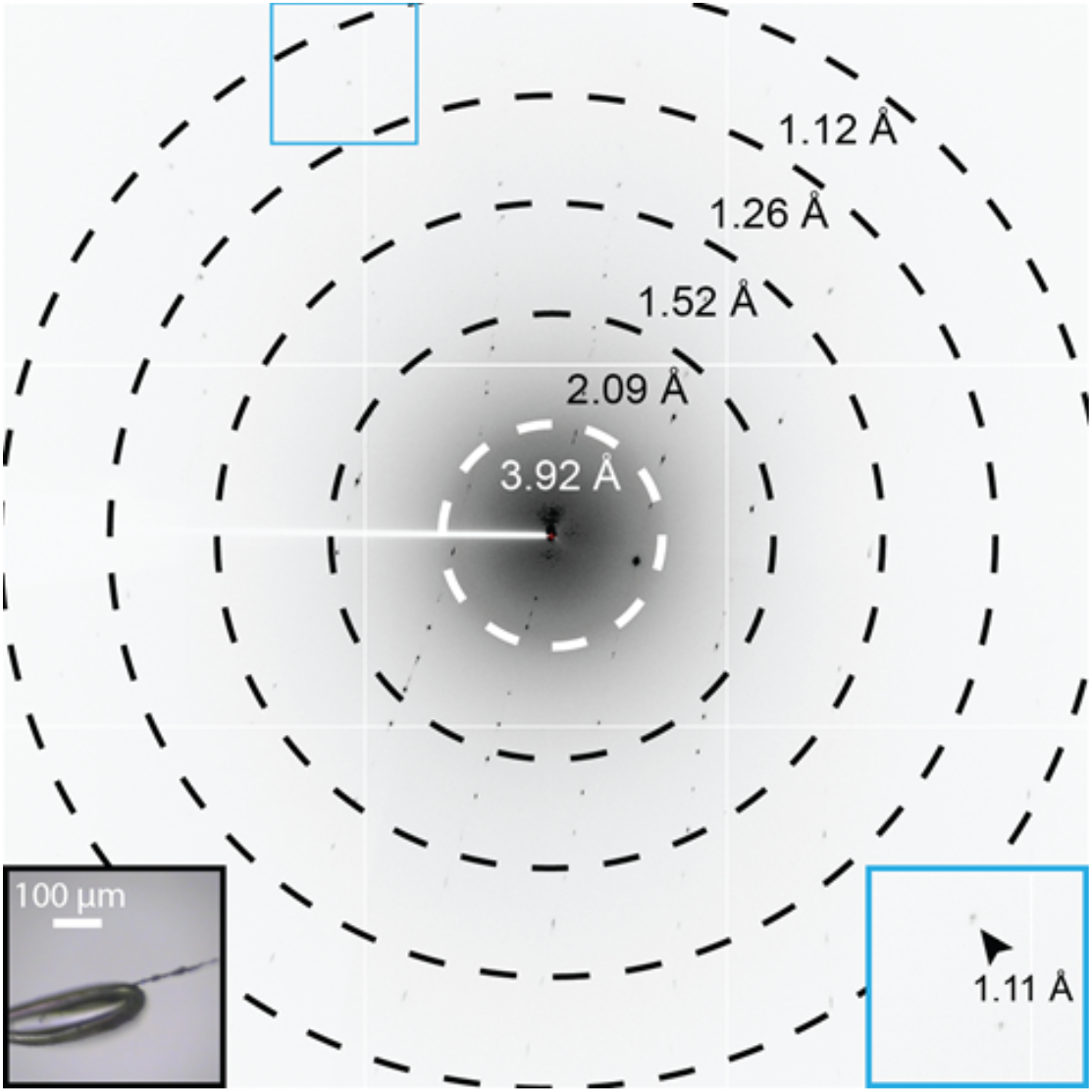
X-ray diffraction of the NYNNYQ peptide Single diffraction pattern of NYNNYQ measured during manual continuous raster scanning, microfocus x-ray data collection. The pattern corresponds to a 5° wedge of reciprocal space. Black inset shows in-line crystal image; the area boxed by the blue square corresponds to the magnified region (blue inset) of the pattern, which shows diffraction near the detector edge at approximately 1.1 Å resolution (black arrow). Resolution rings are indicated.

**Supplementary Figure 3:**
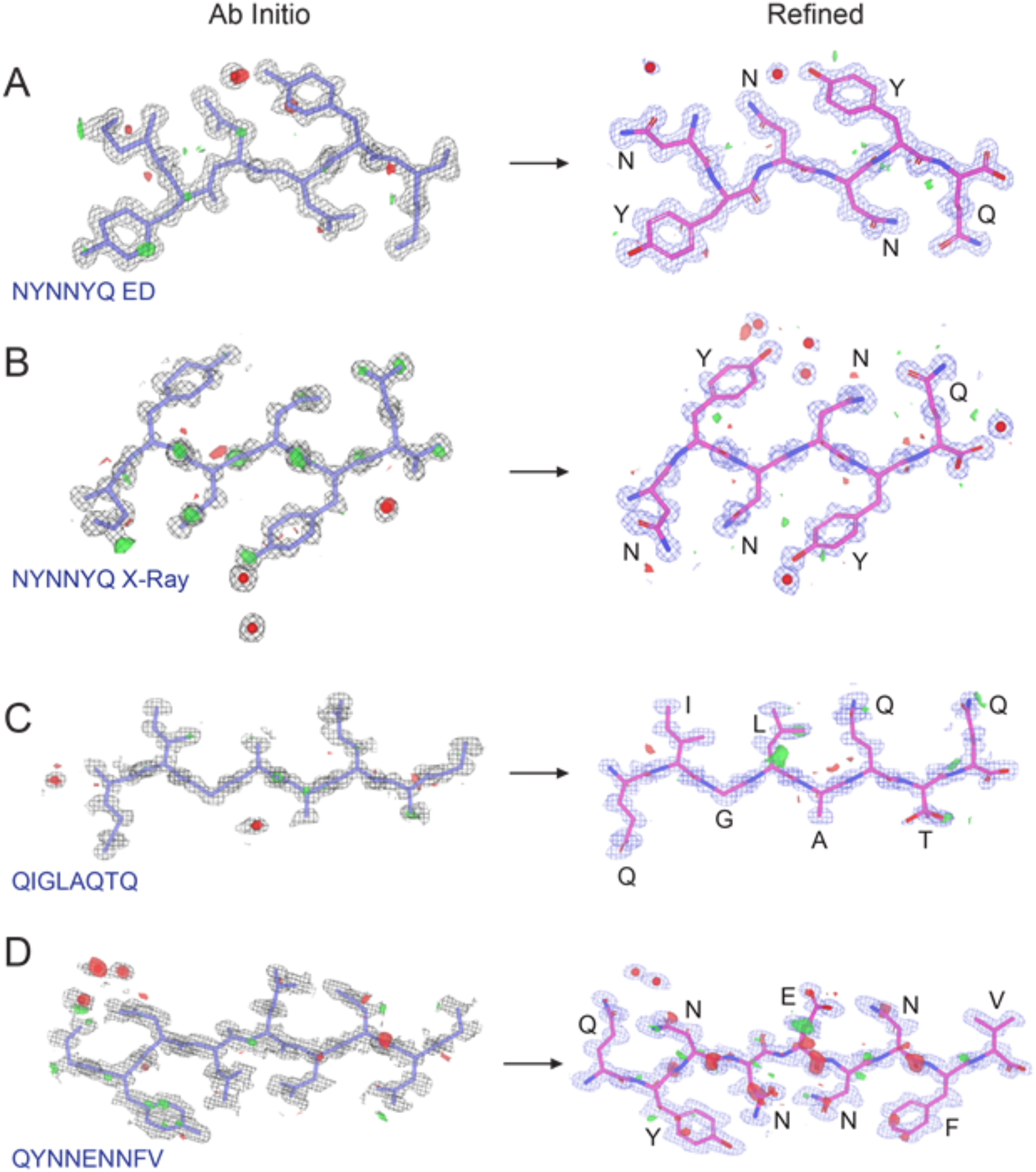
*Ab initio* solutions and refined peptide structures *Ab initio* structures and maps of NYNNYQ (A, electron), NYNNYQ (B, x-ray), QIGLAQTQ (C, plate polymorph), and QYNNENNFV (D). The initial atomic coordinates on the left, calculated by *SHELXD*, are overlaid by their corresponding initial maps. The refined atomic structures on the right are overlaid by their corresponding final maps. The 2F_O_-F_C_ maps, represented by the black mesh (left) or blue mesh (right), is contoured at 1.2σ. Green and red surfaces represent F_o_-F_c_ maps contoured at 3.0σ and -3.0σ, respectively. Modeled waters are present as red spheres. The waters in the *ab initio* solution of B and the refined structure are related by symmetry. The waters in the *ab initio* solution of C were later determined to be backbone atoms during refinement.

**Supplementary Figure 4:**
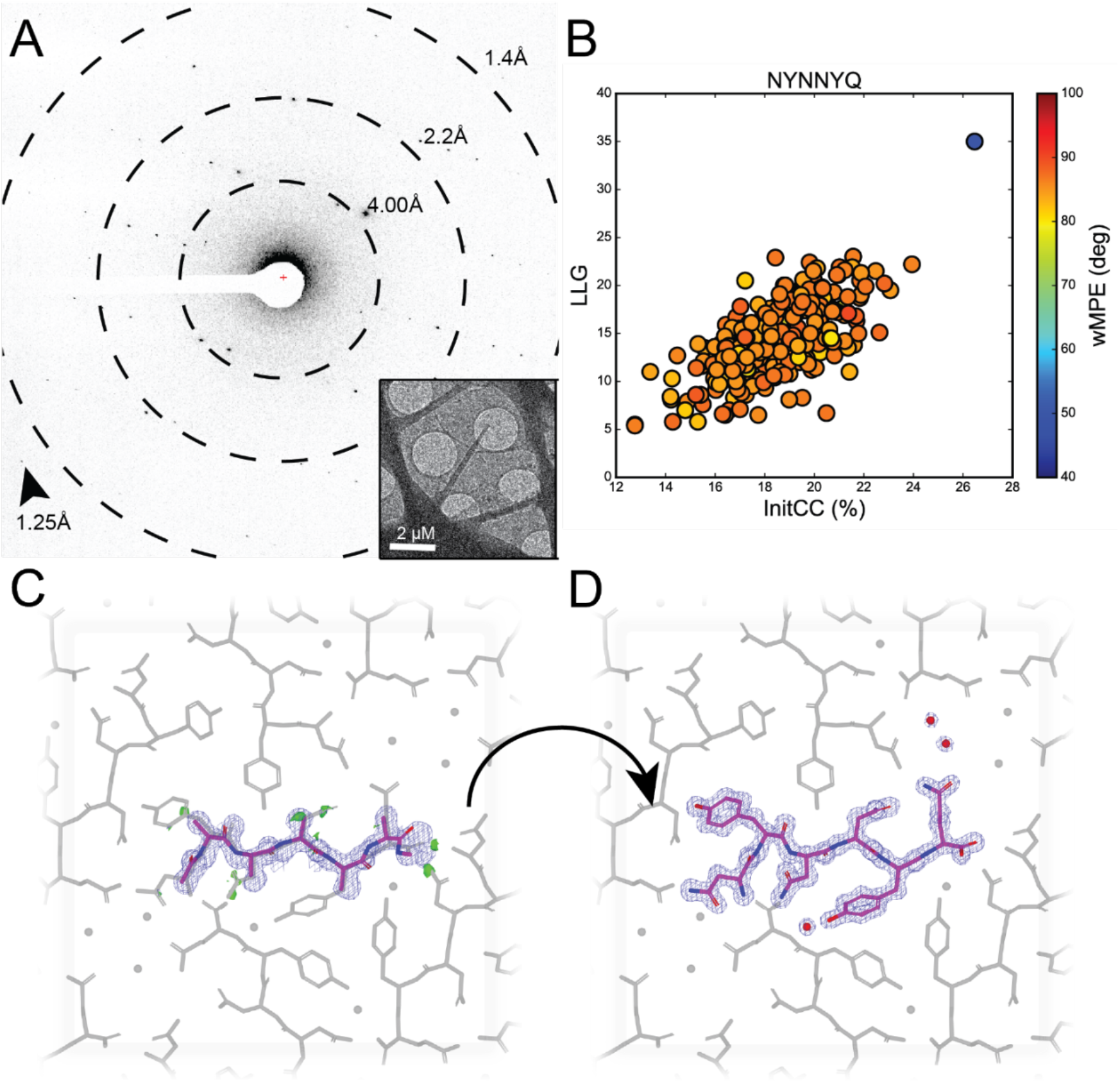
Diffraction pattern, overfocused diffraction image, density maps of initial ARCIMBOLDO outputs and final structures for New1p peptide. A) Diffraction pattern for NYNNYQ B) Post-mortem analysis for ARCIMBOLDO-BORGES plotting LLG vs. InitCC. C) Initial output potential maps expanded by SHELXE for NYNNYQ overlaid on final solution (grey). D) Final potential maps of NYNNYQ with symmetry mates shown in grey.

## Notes

### Competing Interest Statement

JAR is a founder and equity shareholder of Medstruc Inc.

## Literature cited

1. Brink, C. et al. Structure of Vitamin B12: X-ray Crystallographic Evidence on the Structure of Vitamin B12. Nature 174, 1169–1171 (1954).

2. Sawaya, M. R. et al. Ab initio structure determination from prion nanocrystals at atomic resolution by MicroED. Proc. Natl. Acad. Sci. 113, 11232–11236 (2016).

3. Rodriguez, J. A. et al. Structure of the toxic core of α-synuclein from invisible crystals. Nature 525, 486–490 (2015).

4. Gallagher-Jones, M. et al. Sub-ångström cryo-EM structure of a prion protofibril reveals a polar clasp. Nat. Struct. Mol. Biol. 25, 131–134 (2018).

5. Henderson, R. The potential and limitations of neutrons, electrons and X-rays for atomic resolution microscopy of unstained biological molecules. Q. Rev. Biophys. 28, 171–193 (1995).

6. Warmack, R. A. et al. Structure of amyloid-β (20-34) with Alzheimer’s-associated isomerization at Asp23 reveals a distinct protofilament interface. Nat. Commun. 10, 1–12 (2019).

7. Jones, C. G. et al. The CryoEM Method MicroED as a Powerful Tool for Small Molecule Structure Determination. ACS Cent. Sci. 4, 1587–1592 (2018).

8. Ting, C. P. et al. Use of a Scaffold Peptide in the Biosynthesis of Amino Acid Derived Natural Products. Science 365, 280–284 (2019).

9. Sheldrick, G. M. et al. Ab initio phasing. urn:isbn:978-0-470-66078-2 https://it.iucr.org/Fb/ch16o1v0001/ (2012) doi:10.1107/97809553602060000850.

10. Rossmann, M. G. The molecular replacement method. Acta Crystallogr. A 46 (Pt 2), 73–82 (1990).

11. Shi, D., Nannenga, B. L., Iadanza, M. G. & Gonen, T. Three-dimensional electron crystallography of protein microcrystals. eLife 2, e01345 (2013).

12. Krotee, P. et al. Atomic structures of fibrillar segments of hIAPP suggest tightly mated β-sheets are important for cytotoxicity. eLife 6, e19273 (2017).

13. de la Cruz, M. J. et al. Atomic resolution structures from fragmented protein crystals by the cryoEM method MicroED. Nat. Methods 14, 399–402 (2017).

14. Rodríguez, D. D. et al. Crystallographic ab initio protein structure solution below atomic resolution. Nat. Methods 6, 651–653 (2009).

15. Read, R. J. & McCoy, A. J. A log-likelihood-gain intensity target for crystallographic phasing that accounts for experimental error. Acta Crystallogr. Sect. Struct. Biol. 72, 375–387 (2016).

16. Usón, I. & Sheldrick, G. M. An introduction to experimental phasing of macromolecules illustrated by SHELX; new autotracing features. Acta Crystallogr. Sect. Struct. Biol. 74, 106–116 (2018).

17. Millán, C. et al. Combining phase information in reciprocal space for molecular replacement with partial models. Acta Crystallogr. D Biol. Crystallogr. 71, 1931–1945 (2015).

18. Sammito, M. et al. ARCIMBOLDO_LITE: single-workstation implementation and use. Acta Crystallogr. D Biol. Crystallogr. 71, 1921–1930 (2015).

19. Sammito, M. et al. Exploiting tertiary structure through local folds for crystallographic phasing. Nat. Methods 10, 1099–1101 (2013).

20. Millán, C. et al. Exploiting distant homologues for phasing through the generation of compact fragments, local fold refinement and partial solution combination. Acta Crystallogr. Sect. Struct. Biol. 74, 290–304 (2018).

21. Richards, L. S. et al. Fragment-based determination of a proteinase K structure from MicroED data using ARCIMBOLDO_SHREDDER. Acta Crystallogr. Sect. Struct. Biol. 76, 703–712 (2020).

22. Medina, A. et al. ALEPH: a network-oriented approach for the generation of fragment-based libraries and for structure interpretation. Acta Crystallogr. Sect. Struct. Biol. 76, 193–208 (2020).

23. Millán, C., Jiménez, E., Schuster, A., Diederichs, K. & Usón, I. ALIXE: a phase-combination tool for fragment-based molecular replacement. Acta Crystallogr. Sect. Struct. Biol. 76, 209–220 (2020).

24. Thorn, A. & Sheldrick, G. M. Extending molecular-replacement solutions with SHELXE. Acta Crystallogr. D Biol. Crystallogr. 69, 2251–2256 (2013).

25. Alford, R. F. et al. The Rosetta all-atom energy function for macromolecular modeling and design. J. Chem. Theory Comput. 13, 3031–3048 (2017).

26. Chaudhury, S., Lyskov, S. & Gray, J. J. PyRosetta: a script-based interface for implementing molecular modeling algorithms using Rosetta. Bioinforma. Oxf. Engl. 26, 689–691 (2010).

27. Goldschmidt, L., Teng, P. K., Riek, R. & Eisenberg, D. Identifying the amylome, proteins capable of forming amyloid-like fibrils. Proc. Natl. Acad. Sci. 107, 3487–3492 (2010).

28. Caballero, I. et al. ARCIMBOLDO on coiled coils. Acta Crystallogr. Sect. Struct. Biol. 74, 194–204 (2018).

29. Tayeb-Fligelman, E. et al. The cytotoxic Staphylococcus aureus PSMα3 reveals a cross-α amyloid-like fibril. Science 355, 831–833 (2017).

30. Salinas, N. et al. The amphibian antimicrobial peptide uperin 3.5 is a cross-α/cross-β chameleon functional amyloid. Proc. Natl. Acad. Sci. 118, (2021).

31. Senior, A. W. et al. Improved protein structure prediction using potentials from deep learning. Nature 577, 706–710 (2020).

32. Kabsch, W. Integration, scaling, space-group assignment and post-refinement. Acta Crystallogr. D Biol. Crystallogr. 66, 133–144 (2010).

33. Usón, I. & Sheldrick, G. M. Advances in direct methods for protein crystallography. Curr. Opin. Struct. Biol. 9, 643–648 (1999).

34. Sheldrick, G. M. Crystal structure refinement with SHELXL. Acta Crystallogr. Sect. C Struct. Chem. 71, 3–8 (2015).

35. McCoy, A. J. et al. Phaser crystallographic software. J. Appl. Crystallogr. 40, 658–674 (2007).

36. Fujinaga, M. & Read, R. J. Experiences with a new translation-function program. J. Appl. Crystallogr. 20, 517–521 (1987).

37. Leaver-Fay, A. et al. Rosetta3: An Object-Oriented Software Suite for the Simulation and Design of Macromolecules. Methods Enzymol. 487, 545–574 (2011).

38. Liebschner, D. et al. Macromolecular structure determination using X-rays, neutrons and electrons: recent developments in Phenix. Acta Crystallogr. Sect. Struct. Biol. 75, 861–877 (2019).

39. Afonine, P. V. et al. Towards automated crystallographic structure refinement with phenix.refine. Acta Crystallogr. D Biol. Crystallogr. 68, 352–367 (2012).

40. Emsley, P., Lohkamp, B., Scott, W. G. & Cowtan, K. Features and development of Coot. Acta Crystallogr. D Biol. Crystallogr. 66, 486–501 (2010).

## Supplementary Citations

1. Apostol, M. I., Wiltzius, J. J. W., Sawaya, M. R., Cascio, D. & Eisenberg, D. Atomic Structures Suggest Determinants of Transmission Barriers in Mammalian Prion Disease. Biochemistry 50, 2456–2463 (2011).

2. Brumshtein, B. et al. Identification of two principal amyloid-driving segments in variable domains of Ig light chains in systemic light-chain amyloidosis. J. Biol. Chem. 293, 19659–19671 (2018).

3. Colletier, J.-P. et al. Molecular basis for amyloid-polymorphism. Proceedings of the National Academy of Sciences 108, 16938–16943 (2011).

4. Colvin, M. T. et al. Atomic Resolution Structure of Monomorphic Aβ 42 Amyloid Fibrils. J. Am. Chem. Soc. 138, 9663–9674 (2016).

5. Do, T. D. et al. Factors That Drive Peptide Assembly from Native to Amyloid Structures: Experimental and Theoretical Analysis of [Leu-5]-Enkephalin Mutants. J. Phys. Chem. B 118, 7247–7256 (2014).

6. Evans, K. C., Berger, E. P., Cho, C. G., Weisgraber, K. H. & Lansbury, P. T. Apolipoprotein E is a kinetic but not a thermodynamic inhibitor of amyloid formation: implications for the pathogenesis and treatment of Alzheimer disease. Proceedings of the National Academy of Sciences 92, 763–767 (1995).

7. Fitzpatrick, A. W. P. et al. Cryo-EM structures of tau filaments from Alzheimer’s disease. Nature 547, 185–190 (2017).

8. Gremer, L. et al. Fibril structure of amyloid-β(1–42) by cryo–electron microscopy. Science 358, 116–119 (2017).

9. Guenther, E. L. et al. Atomic structures of TDP-43 LCD segments and insights into reversible or pathogenic aggregation. Nat Struct Mol Biol 25, 463–471 (2018).

10. Hughes, M. P. et al. Atomic structures of low-complexity protein segments reveal kinked β sheets that assemble networks. Science 359, 698–701 (2018).

11. Ivanova, M. I. et al. Aggregation-triggering segments of SOD1 fibril formation support a common pathway for familial and sporadic ALS. Proceedings of the National Academy of Sciences 111, 197–201 (2014).

13. Laganowsky, A. et al. Atomic View of a Toxic Amyloid Small Oligomer. Science 335, 1228–1231 (2012).

14. Li, D. et al. Structure-Based Design of Functional Amyloid Materials. J. Am. Chem. Soc. 136, 18044–18051 (2014).

15. Liu, J., Doty, T., Gibson, B. & Heyer, W.-D. Human BRCA2 protein promotes RAD51 filament formation on RPA-covered single-stranded DNA. Nat Struct Mol Biol 17, 1260–1262 (2010).

16. Lu, J.-X. et al. Molecular Structure of β-Amyloid Fibrils in Alzheimer’s Disease Brain Tissue. Cell 154, 1257–1268 (2013).

17. Luo, F. et al. Atomic structures of FUS LC domain segments reveal bases for reversible amyloid fibril formation. Nat Struct Mol Biol 25, 341–346 (2018).

18. Murray, D. T. et al. Structure of FUS Protein Fibrils and Its Relevance to Self-Assembly and Phase Separation of Low-Complexity Domains. Cell 171, 615-627.e16 (2017).

19. Nelson, R. et al. Structure of the cross-β spine of amyloid-like fibrils. Nature 435, 773–778 (2005).

20. Rodriguez, J. A. et al. Structure of the toxic core of α-synuclein from invisible crystals. Nature 525, 486–490 (2015).

21. Saelices, L. et al. Amyloid seeding of transthyretin by ex vivo cardiac fibrils and its inhibition. Proc Natl Acad Sci USA 115, E6741–E6750 (2018).

22. Saelices, L. et al. A pair of peptides inhibits seeding of the hormone transporter transthyretin into amyloid fibrils. J. Biol. Chem. 294, 6130–6141 (2019).

23. Sangwan, S., Sawaya, M. R., Murray, K. A., Hughes, M. P. & Eisenberg, D. S. Atomic structures of corkscrew-forming segments of SOD1 reveal varied oligomer conformations: Structures of Variants of a Toxic Segment from SOD1. Protein Science 27, 1231–1242 (2018).

24. Sawaya, M. R. et al. Atomic structures of amyloid cross-β spines reveal varied steric zippers. Nature 447, 453–457 (2007).

25. Schütz, A. K. et al. Atomic-Resolution Three-Dimensional Structure of Amyloid β Fibrils Bearing the Osaka Mutation. Angew. Chem. Int. Ed. 54, 331–335 (2015).

26. Seidler, P. M. et al. Structure-based inhibitors of tau aggregation. Nature Chem 10, 170–176 (2018).

27. Sgourakis, N. G., Yau, W.-M. & Qiang, W. Modeling an In-Register, Parallel “Iowa” Aβ Fibril Structure Using Solid-State NMR Data from Labeled Samples with Rosetta. Structure 23, 216–227 (2015).

28. Sievers, S. A. et al. Structure-based design of non-natural amino-acid inhibitors of amyloid fibril formation. Nature 475, 96–100 (2011).

29. Soragni, A. et al. A Designed Inhibitor of p53 Aggregation Rescues p53 Tumor Suppression in Ovarian Carcinomas. Cancer Cell 29, 90–103 (2016).

30. Soragni, A. et al. Toxicity of Eosinophil MBP Is Repressed by Intracellular Crystallization and Promoted by Extracellular Aggregation. Molecular Cell 57, 1011–1021 (2015).

31. Soriaga, A. B., Sangwan, S., Macdonald, R., Sawaya, M. R. & Eisenberg, D. Crystal Structures of IAPP Amyloidogenic Segments Reveal a Novel Packing Motif of Out-of-Register Beta Sheets. J. Phys. Chem. B 120, 5810–5816 (2016).

32. Tuttle, M. D. et al. Solid-state NMR structure of a pathogenic fibril of full-length human α-synuclein. Nat Struct Mol Biol 23, 409–415 (2016).

33. Wiltzius, J. J. W. et al. Molecular mechanisms for protein-encoded inheritance. Nat Struct Mol Biol 16, 973–978 (2009).

34. Wiltzius, J. J. W. et al. Atomic structure of the cross-β spine of islet amyloid polypeptide (amylin). Protein Sci. 17, 1467–1474 (2008).

35. Xiao, Y. et al. Aβ(1–42) fibril structure illuminates self-recognition and replication of amyloid in Alzheimer’s disease. Nat Struct Mol Biol 22, 499–505 (2015).

36. Yu, L., Lee, S.-J. & Yee, V. C. Crystal Structures of Polymorphic Prion Protein β1 Peptides Reveal Variable Steric Zipper Conformations. Biochemistry 54, 3640–3648 (2015).

